# Intersubject MVPD: Empirical Comparison of fMRI Denoising Methods for Connectivity Analysis

**DOI:** 10.1101/456970

**Authors:** Yichen Li, Rebecca Saxe, Stefano Anzellotti

## Abstract

Noise is a major challenge for the analysis of fMRI data in general and for connectivity analyses in particular. As researchers develop increasingly sophisticated tools to model statistical dependence between the fMRI signal in different brain regions, there is a risk that these models may increasingly capture artifactual relationships between regions, that are the result of noise. Thus, choosing optimal denoising methods is a crucial step to maximize the accuracy and reproducibility of connectivity models. Most comparisons between denoising methods require knowledge of the ground truth: of what is the ‘real signal’. For this reason, they are usually based on simulated fMRI data. However, simulated data may not match the statistical properties of real data, limiting the generalizability of the conclusions. In this article, we propose an approach to evaluate denoising methods using real (non-simulated) fMRI data. First, we introduce an intersubject version of multivariate pattern dependence (iMVPD) that computes the statistical dependence between a brain region in one participant, and another brain region in a different participant. iMVPD has the following advantages: 1) it is multivariate, 2) it trains and tests models on independent folds of the real fMRI data, and 3) it generates predictions that are both between subjects and between regions. Since whole-brain sources of noise are more strongly correlated within subject than between subjects, we can use the difference between standard MVPD and iMVPD as a ‘discrepancy metric’ to evaluate denoising techniques (where more effective techniques should yield smaller differences). As predicted, the difference is the greatest in the absence of denoising methods. Furthermore, a combination of removal of the global signal and CompCorr optimizes denoising (among the set of denoising options tested).

## 1 Introduction

Cognitive tasks elicit the activation of multiple brain regions (Ishai, 2008; Anzellotti and Caramazza, 2015). To understand how these regions function jointly to implement cognition, we need to investigate their connectivity. Connectivity measures based on structural data (i.e. Assaf and Pasternak (2008)) and connectivity measures based on functional data (i.e. Biswal et al. (1995)) have distinct advantages: the former enable inferences about whether regions are directly connected, the latter have the potential to investigate task-specific changes in the interactions between regions. Perhaps the most popular method to study interactions using functional data is ‘functional connectivity’, an analysis technique based on computing Pearson’s correlation between the average responses in two brain regions over time (Biswal et al., 1995). Functional connectivity has been used extensively to map brain networks (Thomas Yeo et al., 2011) and search for biomarkers for patient populations (Drysdale et al., 2017).

Functional connectivity has served as the backbone of a vast literature, but the development of new methods for the study of neural responses and the emergence of new questions called for a new approach to study functional interactions between brain regions. Functional connectivity is univariate in nature: it is based on correlations between spatially-averaged responses. By contrast, it is now known that multivariate patterns of response encode rich information that is lost by averaging (Haxby et al., 2001; Haynes and Rees, 2006). Furthermore, functional connectivity does not lead to a predictive model that can be tested in independent data. Training a model with part of the data and testing its accuracy in independent data has become the norm for some types of analyses (i.e. MultiVoxel Pattern Analysis - MVPA, Norman et al. (2006)), but it is not done in functional connectivity, and this makes it more susceptible to noise. A technique was needed to capture interactions between brain regions preserving multivariate information, and to implement training and testing in independent data. To satisfy these requirements, we have recently developed MultiVariate Pattern Dependence (MVPD, Anzellotti et al. (2017)), and we have found that it is more sensitive than univariate connectivity techniques (Anzellotti et al., 2017).

MVPD fits a model with a set of training data and tests it in independent data, therefore it is more robust to noise than methods lacking an independent ‘generalization’ test. However, some sources of noise may still affect MVPD results. For this reason, it is critical to use effective denoising techniques. Estimating the effectiveness of different denoising techniques for connectivity analysis has been typically challenging. Studies comparing different denoising approaches usually rely on simulations (i.e. Power et al. (2015)), because in simulations the ground truth is known. The simulation model determines what part of the overall response is the signal and what part is the noise, and the failure or success of the denoising methods can be determined unambiguously.

Unfortunately, applying conclusions obtained with simulation models to real fMRI data is non-trivial. Despite efforts to generate simulations that preserve many of the statistical properties of Blood-Oxygen Level Dependent (BOLD) signal, it is very difficult to generate simulations that perfectly match the statistical properties of real BOLD timecourses. This difficulty poses a major obstacle towards developing increasingly sophisticated connectivity techniques: more complex connectivity models might capture complex relationships between the noise in different regions, and thus require increasingly effective denoising. The differences between simulations and real data could thus have an even greater impact on the evaluation of denoising techniques.

In this article, we developed an approach to evaluate different denoising techniques using real fMRI data (as opposed to simulations). Our approach relies on computing interactions between brain regions between participants, following an ‘intersubject’ approach. Intersubject versions of univariate connectivity measures have been used in the previous literature to study individual differences in the responses to the same stimuli (intersubject correlations, Hasson et al. (2004, 2008); Wilson et al. (2007)). We introduce here an intersubject version of MVPD (iMVPD), which predicts responses between brain regions in different participants (source code available at https://github.com/yl3506/iMVPD_denoise). Three key properties define iMVPD: 1) it is multivariate, 2) it trains and tests models on independent folds of the data, and 3) it generates predictions that are both between subjects and between regions.

We took advantage of the unique properties of iMVPD to test the effectiveness of different denoising techniques on real fMRI data in the context of multivariate connectivity analyses. Considering a set of functionally-defined regions of interest, we computed connectivity matrices showing MVPD strength across each pair of regions both within subject and between subjects. Since different participants are likely to exhibit variation in their head movements and patterns of breathing or respiration, iMVPD should be less affected by these sources of noise. As a consequence, effective denoising techniques should yield a reduced difference between the within-subject and intersubject connectivity matrices. This gave us an index that we could use to compare different denoising approaches.

## 2 Methods

### 2.1 Data

All analyses in this project use the 3 tesla (3T) audio-visual fMRI dataset from the StudyForrest project (http://studyforrest.org), which encompasses over 2 hours of scans for each of 15 participants (all right handed, mean age 29.4 years, range 21-39, 6 females). T2*-weighted echo-planar images (TR=2s, TE=30m, 90 deg flip angle, parallel acquisition with sensitivity encoding (SENSE) reduction factor 2) were acquired using a whole-body Philips Achieva dStream MRI scanner with a 32 channel head coil. Each volume consists of 35 axial slices with a 10% inter-slice gap. Each slice comprises 80 × 80 voxels, with voxel resolution = 3 × 3 × 3mm, covering a Field of view (FoV) of 240mm (see (Hanke et al., 2016) for additional details).

The dataset includes localizer sessions with stimuli from multiple categories, and the viewing of the entire movie Forrest Gump, subdivided into 8 sections presented to participants in 8 separate functional runs. This dataset was chosen for this study as it provides complex sensory input that follows the same timecourse between participants (Labs et al., 2015). During the localizer session for this experiment, participants were shown 24 unique grayscale images from each of six stimulus categories: human faces, human bodies without heads, small objects, houses and outdoor scenes comprising of nature and street scenes, and phase scrambled images (Sengupta et al., 2016). During the movie session for this experiment, participants were shown two-hour audio-visual stimuli (the movie Forrest Gump) (Joshi et al., 2018).

### 2.2 Preprocessing

The format and folder structure of the dataset was modified to match the BIDS standard (Gorgolewski et al., 2016), and the data were preprocessed using fMRIPrep (https://fmriprep.readthedocs.io/en/latest/index.html): a preprocessing tool that takes advantage of nipype (https://nipype.readthedocs.io/en/latest/) to combine efficient algorithms for fMRI preprocessing from different software packages, minimizing experimenter degrees of freedom and offering a controlled environment that favors reproducibility (Esteban et al., 2018). Specifically, FSL MCFLIRT (https://fsl.fmrib.ox.ac.uk/fsl/fslwiki/MCFLIRT) was used to estimate head motion, and each individual’s functional data were coregistered to her/his anatomical scan. Segmentation and normalization were performed with ANTs (http://stnava.github.io/ANTs/).

### 2.3 Regions of Interest (ROIs) Definition

In order to make the results easily replicable, we analyzed two networks of category-selective brain regions that are highly reliable between participants and widely studied in the literature (Figure 1): the face-selective network (including the occipital face area - rOFA, fusiform face area - rFFA, anterior temporal lobe - rATL, and superior temporal sulcus - rSTS) and the scene-selective network (including transverse occipital sulcus - rTOS, parahippocampal place area - rPPA, and posterior cingulate - rPC).

**Figure 1:**
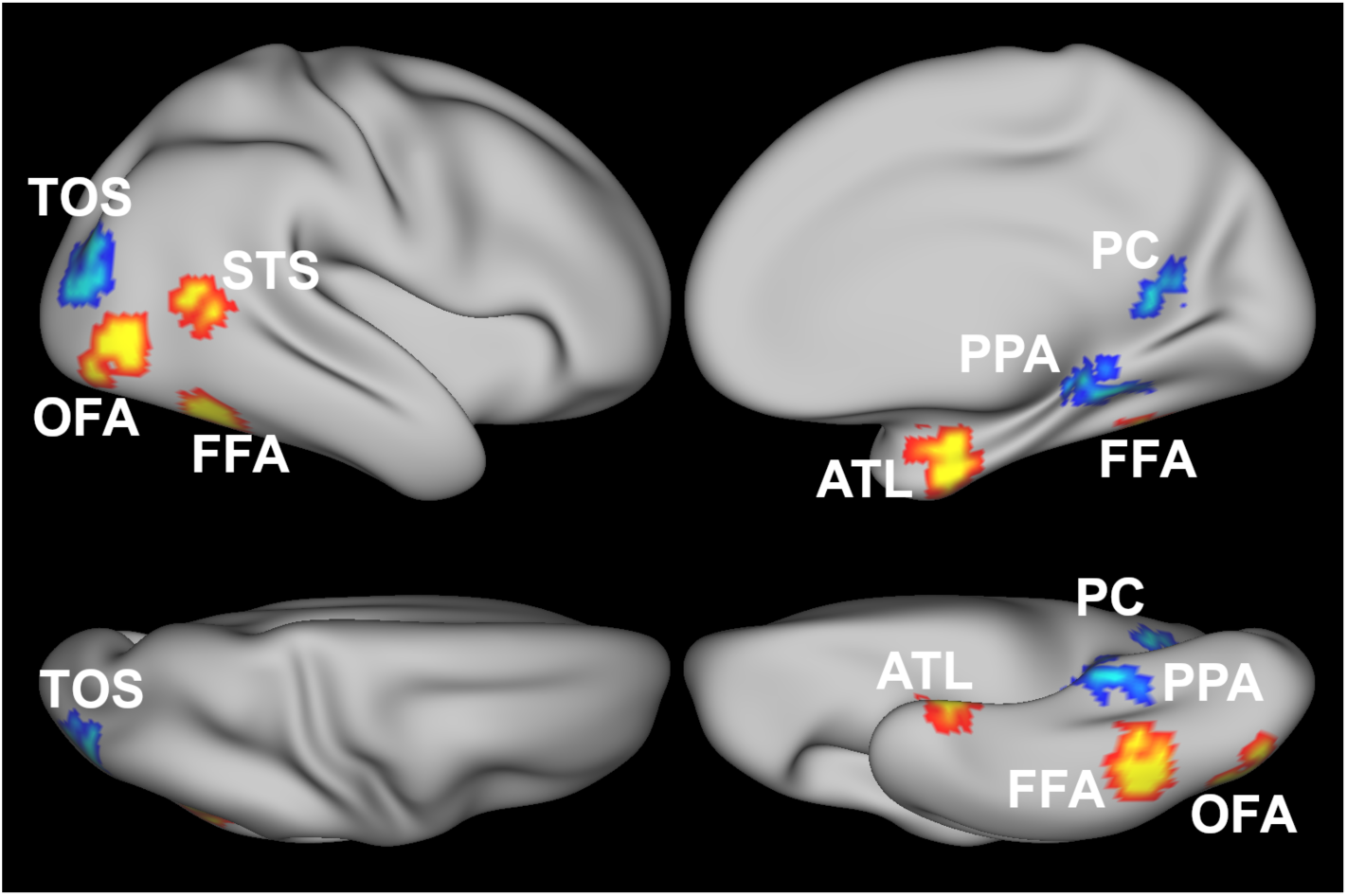
Regions of interest for one example participant shown on an inflated cortical surface. Face-selective regions are shown in red-yellow (occipital face area - OFA, fusiform face area - FFA, superior temporal sulcus - STS, anterior temporal lobe - ATL), scene-selective regions are shown in blue (temporo-occipital sulcus -TOS, parahippocampal place area - PPA, posterior cingulate/retrosplenial cortex - PC).

Given the similarity in the responses in these regions across hemispheres, we restricted our analysis to the right hemisphere. First-level general linear models (GLM) were estimated in each participant with FSL FEAT (https://fsl.fmrib.ox.ac.uk/fsl/fslwiki/FEAT) using the independent localizer data. Face-selective regions were identified in individual participants using the contrast faces *>* scenes in an independent functional localizer. Anatomical parcels generated from a large number of independent subjects were used as a search space to identify contrast peaks for each of the regions. We then defined a 9mm sphere centered in the peak and selected the 80 voxels showing strongest face-selectivity within the sphere, as assessed with the contrast t-map. An analogous procedure was adopted for the scene regions, with the exception that the search space mask was defined by visually determining the group-level peaks for the contrast scenes *>* faces. Regions of interest were visualized on the cortical surface with Connectome Workbench (https://www.humanconnectome.org/software/connectome-workbench, Figure 1).

### 2.4 Denoising Methods

We compared the effectiveness of four different denoising approaches for fMRI data: regression of slow trends (Diedrichsen and Shadmehr, 2005), commonly adopted to remove ‘scanner drift’: low frequency fluctuations attributed to physiological noise and subject motion (Smith et al., 1999); regression of the six motion parameters generated during motion correction (Friston et al., 1996), which attempts to remove noise that is linearly related to translations and rotations of the head; removal of the global signal (Macey et al., 2004), which dicards the variability in a voxel’s responses that is shared with the fluctuation of the average signal in the entire brain; and CompCorr (Behzadi et al., 2007), which extracts principal components from the signal in the white matter and cerebrospinal fluid and regresses them out from each voxel. All denoising methods were implemented by first generating one (i.e. in the case of global signal) or more (i.e. in the case of motion parameters) predictors. The predictors were then regressing out of each voxel’s responses, and the residuals of the regression were used as the ‘denoised’ signal. Translation and rotation predictors and predictors for the removal of slow trends were obtained from the fMRIPrep outputs. Predictors for CompCorr were computed by generating an eroded mask of the white matter and cerebrospinal fluid using the segmented anatomical data from individual participants, and extracting 5 principal components. The predictor for the global signal was computed averaging the responses in all voxels in a subject-specific gray matter mask generated by fMRIPrep during segmentation In addition to testing the effectiveness of denoising methods taken individually, we assessed combinations of multiple denoising methods to identify optimal noise removal approaches. In total, we compared 11 different denoising pipelines.

### 2.5 iMVPD: Modeling Representational Spaces

For both within subject MVPD, and between subject iMVPD, the data were divided into a 8 folds; each analysis used 7 folds for training and one for testing, iteratively. Using only the training data, dimensionality reduction was applied to the multivariate responses in each region using PCA. Second, a linear function was estimated to predict the responses along the PCs in the target region from the responses in the predictor region (still using only the training data). Finally, the testing data were projected on the PC dimensions estimated with the training data, and the function *f* estimated with the training data was applied to the testing data in the predictor region to generate a prediction for the testing data in the target region. The proportion of variance explained by this prediction in the observed testing data in the target region was computed and used as a measure of multivariate statistical dependence between the two regions. The key difference between MVPD and between subjects iMVPD was that in iMVPD the target regions’ data were drawn from a different participant.

More specifically, let’s consider two participants *A*, *B* and two brain regions (a ‘predictor’ region in participant *A*, and a ‘target’ region in participant *B*). For participant *A*, we extracted multivariate timecourses of response 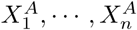 in the predictor region, and for participant *B* we extracted multivariate timecourses 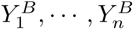 in the target region (where *n* = 8 is the number of runs).

The multivariate timecourse 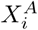 in the predictor region for participant *A* in run *i* was a matrix of size 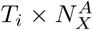, where *T_i_* was the number of time points of the *i^th^* run, and 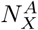 was the number of voxels in the predictor region for participant *A*. Analogously, the multivariate timecourse 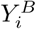 in the target region for participant *B* in run *i* was a matrix of size 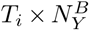,where *T_i_* was the number of time points of the *i^th^* run, and 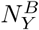 was the number of voxels in the target ROI for participant *B*.

Just as in standard MVPD (Anzellotti et al., 2017), for each choice of a testing run *i*, data in the remaining runs were concatenated, obtaining the training predictor:

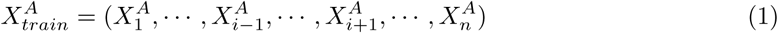

The training target will be obtained by the same procedure, applied to the data from the predictor region from a different subject B:

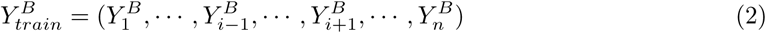

After the generation of the training set, principal component analysis (PCA) was applied to 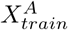 and 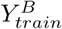:

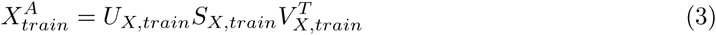

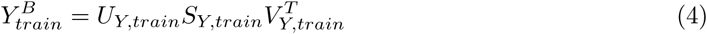

Dimensionality reduction was implemented projecting 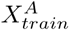 and 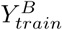 on lower dimensional sub-spaces spanned by the first *k_X_* and *k_Y_* principal components respectively:

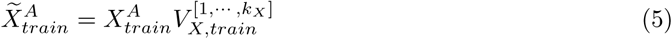

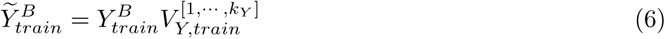

where 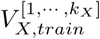 was the matrix formed by the first *k_X_* columns of *V_X,train_*, and 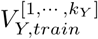 was the matrix formed by the first *k_Y_* columns of *V_Y,train_*. For each region, each dimension obtained with PCA was a linear combination of the voxels in the region, whose weights defined a multivariate pattern of response over voxels. Considering as an example the predictor region, the scores of a dimension *k* were encoded in the *k^th^* column of 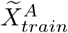, and represented the intensity with which the multivariate pattern corresponding to dimension *k* was activated over time. In previous work, we found that using 3 components outperformed using 1 or 2 components (Anzellotti et al., 2017). To test this observation in this new dataset, we compared the MVPD results obtained from choosing 1, 2, and 3 components using a default denoising method – removal of slow trends + CompCorr (will be explained in section 4). Consistently with prior findings (Anzellotti et al., 2017), the model with 3 components explained the most independent variance (see section 2.7): averaging between all region pairs we obtained the following values of ‘multivariate independent correlation index’ (square root of the independent variance explained): 0.2169, 0.2560, 0.2774 respectively for 1, 2 and 3 principal components. Therefore, we used 3 components for the rest of our analyses.

### 2.6 iMVPD: Modeling Statistical Dependence and Predicting Multivariate Timecourses

The mapping *f* from the dimensionality-reduced timecourses in the predictor region 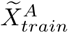 to the dimensionality-reduced timecourses in the target region 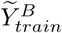 was modeled with multiple linear regression:

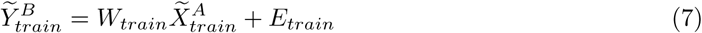

where the model parameters were estimated using ordinary least squares (OLS).

The extension of multivariate analysis methods to intersubject formulations can pose unique challenges. For example, in MVPA, classification between participants is usually poor unless functional alignment techniques are not first used to identify a common response space between participants (Guntupalli and Haxby, 2010; Haxby et al., 2011). In the case of iMVPD, despite the brain region used as predictor and the brain region that is the target of prediction are in different participants, the learned prediction function is always applied to (independent subsets of) data from the same subject during both training and testing. This ensures that the function learned during training can be applied at the testing stage without the need of additional hyperalignment.

After the estimation of parameters *W_train_*, predictions for the multivariate responses in the left out run *i* were computed by 1) projecting the predictor region’s data in the test run *i* on the principal components of the predictor region estimated with the other runs (the training runs), and 2) multiplying the dimensionality-reduced testing data by the parameters estimated using data from the other runs (the training runs). More formally, for each run *i*, we generated dimensionality reduced responses in the predictor region:

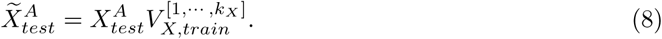

Then, we calculated the predicted responses in the seed region in run *i*:

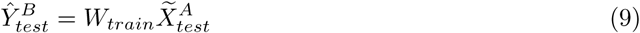

using the parameters *W_train_* independently estimated with the training runs.

### 2.7 Variance explained

To compute the proportion of variance explained in the test data we adopted the following procedure: 1) we applied principal component analysis to the test data, 2) we projected the observed and predicted responses on the principal components obtained from the test data, 3) we calculated the proportion of variance explained by the prediction in each component, and 4) we weighted these proportions by the proportion of variance each component explained in the overall observed response.

More specifically, we calculated

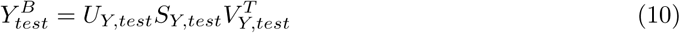

(step 1 above) and we projected the predicted and observed responses on this common orthogonal basis set (step 2):

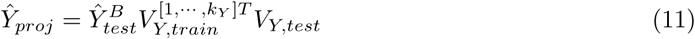

(where 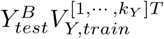 projects the predicted responses from the PCA space computed with the training data to voxel space, and the product by *V_Y,test_* projects them on the PCA space computed with the testing data) and

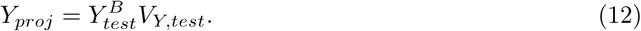

The proportion of variance explained in each component *i* was computed as

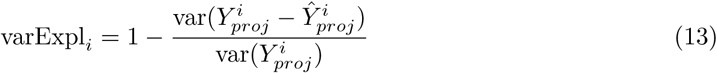

where 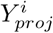 denotes the *i*-th column of matrix *Y_proj_*, which contains the timecourse along component *i* of the PCA space computed with the testing data (step 3). Finally, the total variance explained was calculated as a weighted sum of the variance explained along each dimension (step 4), where the weights are given by the proportion of variance explained by that dimension in the overall response (which can be calculated using the PCA eigenvalues):

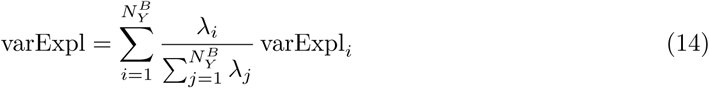

For all analyses involving comparisons between within-subject MVPD and iMVPD, we did not compute predictions from a region to itself (i.e. values on the diagonal) because they are not meaningful for the within-subject analyses, and thus cannot be compared across methods.

### 2.8 Comparing denoising methods

To compare denoising models, we noted that an individual participant’s observed multivariate responses in a region *Y*(*t*) can be decomposed as

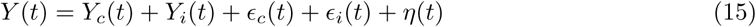

where the term *Y_c_*(*t*) is the part of the true response that is common across individuals, *Y_i_*(*t*) is the part of the true response that is specific to the particular individual, *ϵ_c_* is the whole-brain noise that is shared across individuals and *ϵ_i_*(*t*) is the whole-brain noise that is specific to that individual, and *η*(*t*) is region-specific noise.

A model of connectivity within individual can aspire to explain an amount of variance given by

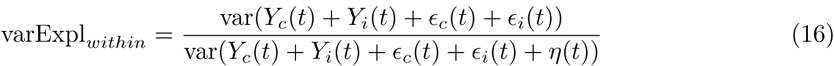

by contrast, an intersubject model can aspire to explain an amount of variance given by

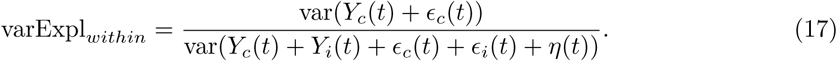

If a denoising method is removing in equal proportions signal and noise, (i.e. by a multiplicative factor 0 *< λ <* 1), we obtain new proportions of variance explained:

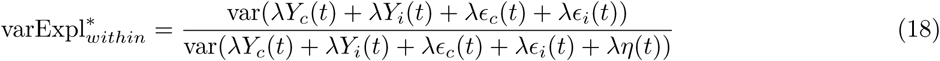

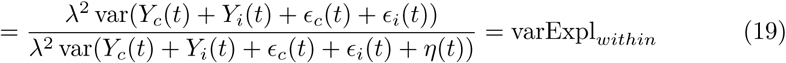

and similarly

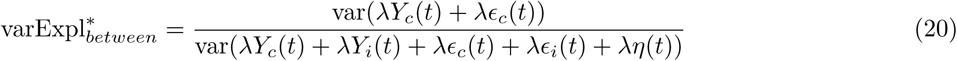

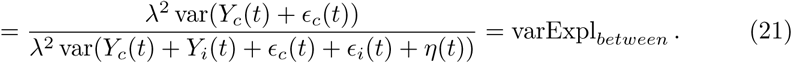

By contrast, if a denoising method disproportionately reduces within-subject whole brain noise, we will have that the numerator of varExpl*_within_* gets proportionally smaller than the numerator of varExpl*_between_*. This is because the term that is disproportionately reduced by denoising is *ϵ_i_*(*t*), which only appears in the numerator of varExpl*_within_*. This effectively reduces the difference

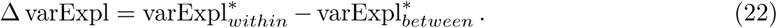

Relying on this logic, we compared the alternative denoising methods using the index Δ varExpl, where values closer to zero indicate better denoising performance. We computed the index Δ varExpl for each pair of brain regions, obtaining ‘connectivity-matrix-like’ figures in which each cell depicts the Δ varExpl index for the corresponding connection.

The average Δ varExpl index across all regions provides a measure of the efficacy of a denoising method (lower is better). Furthermore, we compared the pattern of denoising between regions for different denoising methods calculating the correlation between Δ varExpl matrices, obtaining a measure of similarity between denoising methods in terms of the set of connections on which they are more or less effective.

## 3 Results

### 3.1 Validating Intersubject MVPD

In a first analysis, we tested whether intersubject MVPD is able to identify expected patterns of statistical dependence between different brain regions. Specifically, we tested whether intersubject MVPD, like within-subject MVPD, shows stronger interactions between regions that belong to the same network (i.e. face-selective regions vs scene-selective regions). We found that interactions among regions in the same network were indeed stronger also when using intersubject MVPD, a comparison of connectivity matrices is shown in Figure 2 (see sections 2.4,2.5). To measure the similarity between the patterns of interactions between regions across different region pairs for within-subject and intersubject connectivity we computed the Pearson’s correlation between the connectivity matrices generated with the two methods, obtaining an *r* value of 0.8596. In both within-subject MVPD and iMVPD, strong statistical dependence was found across pairs of region within a same network (defined based on response selectivity in an independent localizer): on average, pairs of face-selective regions showed stronger statistical dependence with other face-selective regions than with scene-selective regions, and viceversa (Figure 2). Relatively accurate predictions were also observed when face-selective regions were used as predictors for some scene-selective regions (in particular the rTOS), but not viceversa. This effect was observed with within-subject and also with intersubject MVPD.

**Figure 2:**
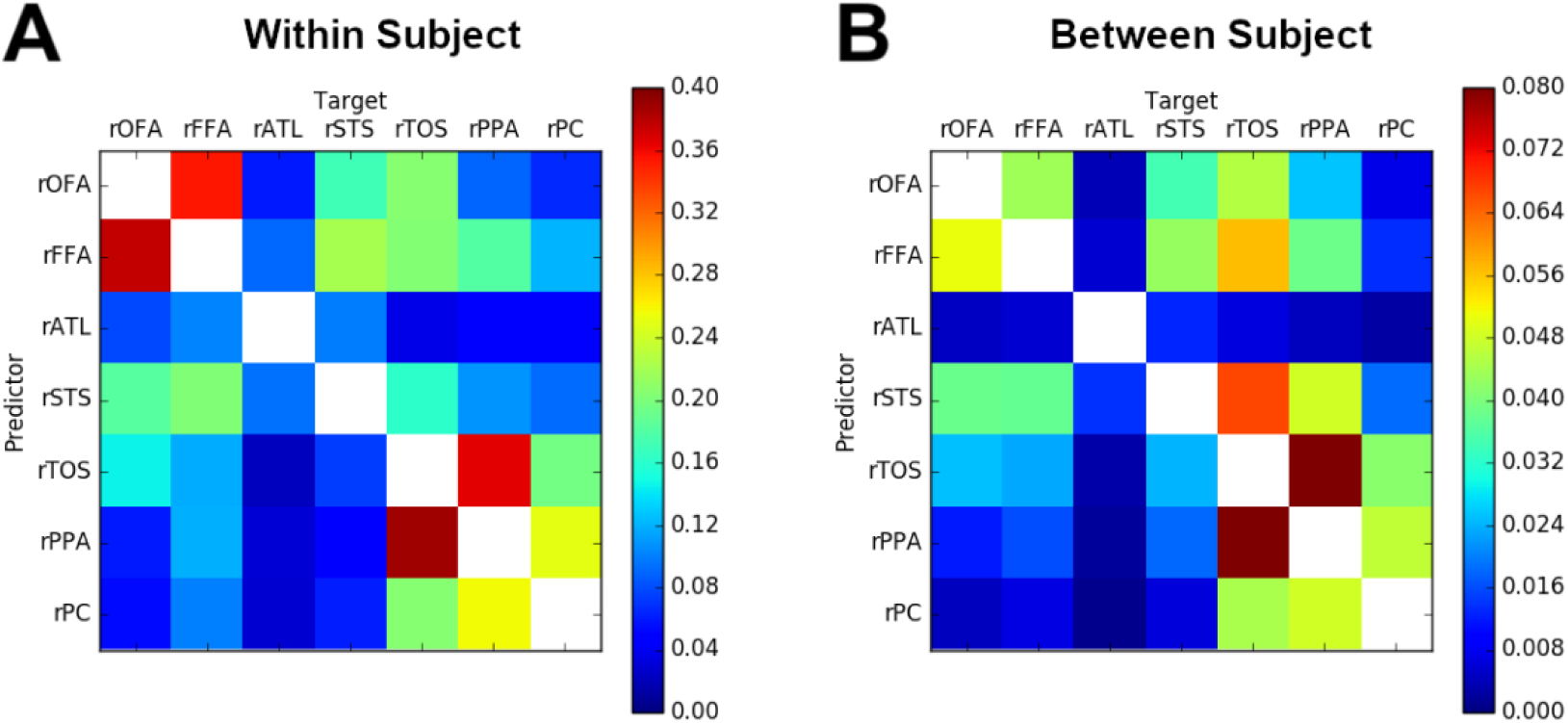
Amount of raw variance explained (as defined in Section 2.7) for within-subject MVPD (A) and between-subject MVPD (B). Each element in a matrix represents the average between participants and cross-validation iterations of the proportion of the variance explained by predicting the multivariate responses in the target region from the the multivariate responses in the predictor region. The similarity between the two matrices was assessed with Pearson’s correlation, yielding *r* = 0.8596.

### 3.2 Single Denoising Methods

After determining that iMVPD yielded similar patterns of statistical dependence across region pairs as within-subject MVPD, we proceeded to assess the efficacy of different denoising approaches (when considered individually) using the difference between the proportion of variance explained in the within-subject analysis minus the proportion of variance explained in the intersubject analysis (Δ varExpl, see Section 2.8).

As a sanity check, we calculated Δ varExpl without applying any denoising method, and indeed we observed a higher value of Δ varExpl than the ones observed after applying any of the other denoising methods (Figure 3A, F). Additionally, we observed differences in the Δ varExpl index across the denoising methods, with CompCorr achieving the best performance (as assessed by Δ varExpl).

**Figure 3:**
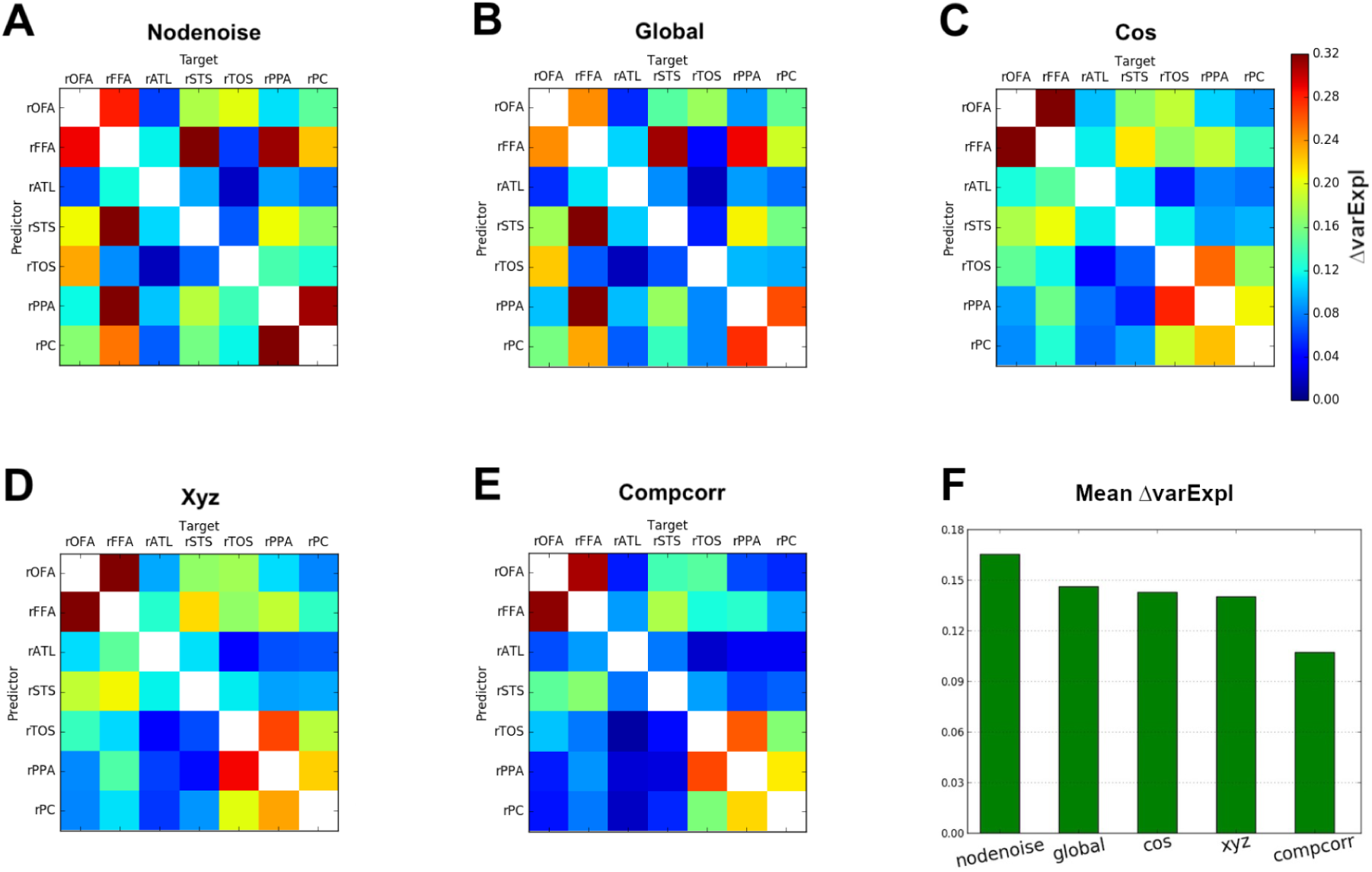
Matrices showing the difference between the variance explained for within-subject MVPD minus the variance explained for iMVPD (Δ varExpl) of A) data without any denoising applied, B) removal of global signal, C) removal of slow trends, D) regression of head translation and rotation parameters (motion parameters), E) Compcorr. Figure F) shows the mean Δ varExpl of the previous five denoising methods in descending order.

Different denoising methods varied not only in terms of the average value of Δ varExpl, but also in the pattern of Δ varExpl across different region pairs. In order to perform a quantitative evaluation of the similarity between denoising methods in terms of their pattern of Δ varExpl across region pairs (that is, in terms of whether they removed noise from similar or distinct sets of connections), we calculated the Pearson’s correlation between the Δ varExpl matrices obtained for the different methods. This analysis revealed largely similar patterns for the removal of slow trends and the regression of translation and rotation parameters. The pattern of noise removal for CompCorr was also relatively similar, but the pattern for removal of global signal was markedly distinct, affecting different pairs of brain regions (Figure 4).

**Figure 4:**
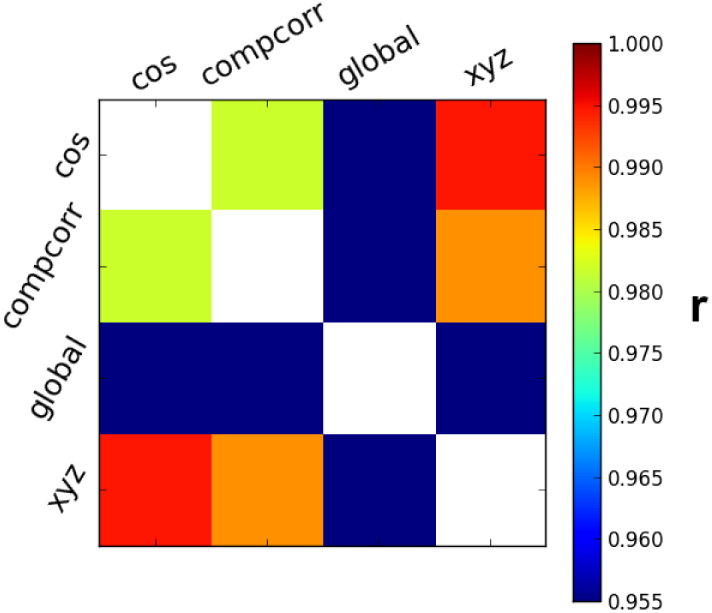
Pearson’s correlation (*r*) of the Δ varExpl across all region pairs of the single denoising methods showed in Figure 3.

### 3.3 Comparing Combinations of Denoising Methods

Multiple denoising methods can be used jointly to improve the efficacy of noise-removal. However, using too many denoising predictors may lead to the risk of removing meaningful variability in the signal. Therefore, we aimed to test different combinations of denoising methods to identify a minimal and effective noise-removal procedure which reduces Δ varExpl without including unnecessary predictors.

To this end, we compared several combinations of denoising methods. Since the removal of slow trends is widely used as a denoising method, we investigated the performance obtained by combining it with the other methods we assessed in the previous analyses (Figure 5 A-C). First, we observed that adding motion regressors (translation and rotation) did not improve denoising appreciably (compare Figure 5A to Figure 3C). Note that this is not a trivial consequence of the high correlation between the Δ varExpl matrices for these two methods, because distinct denoising methods might remove non-overlapping variance in similar amounts from the same set of brain regions, thus yielding high correlations between Δ varExpl matrices without redundancy in their denoising contribution. For example, two different denoising methods might remove two noise sources that are independent of each other in the timecourse of response within a region, but these independent noise sources might nonetheless have a similar distribution in terms of the proportion of noise they generate across region pairs.

**Figure 5:**
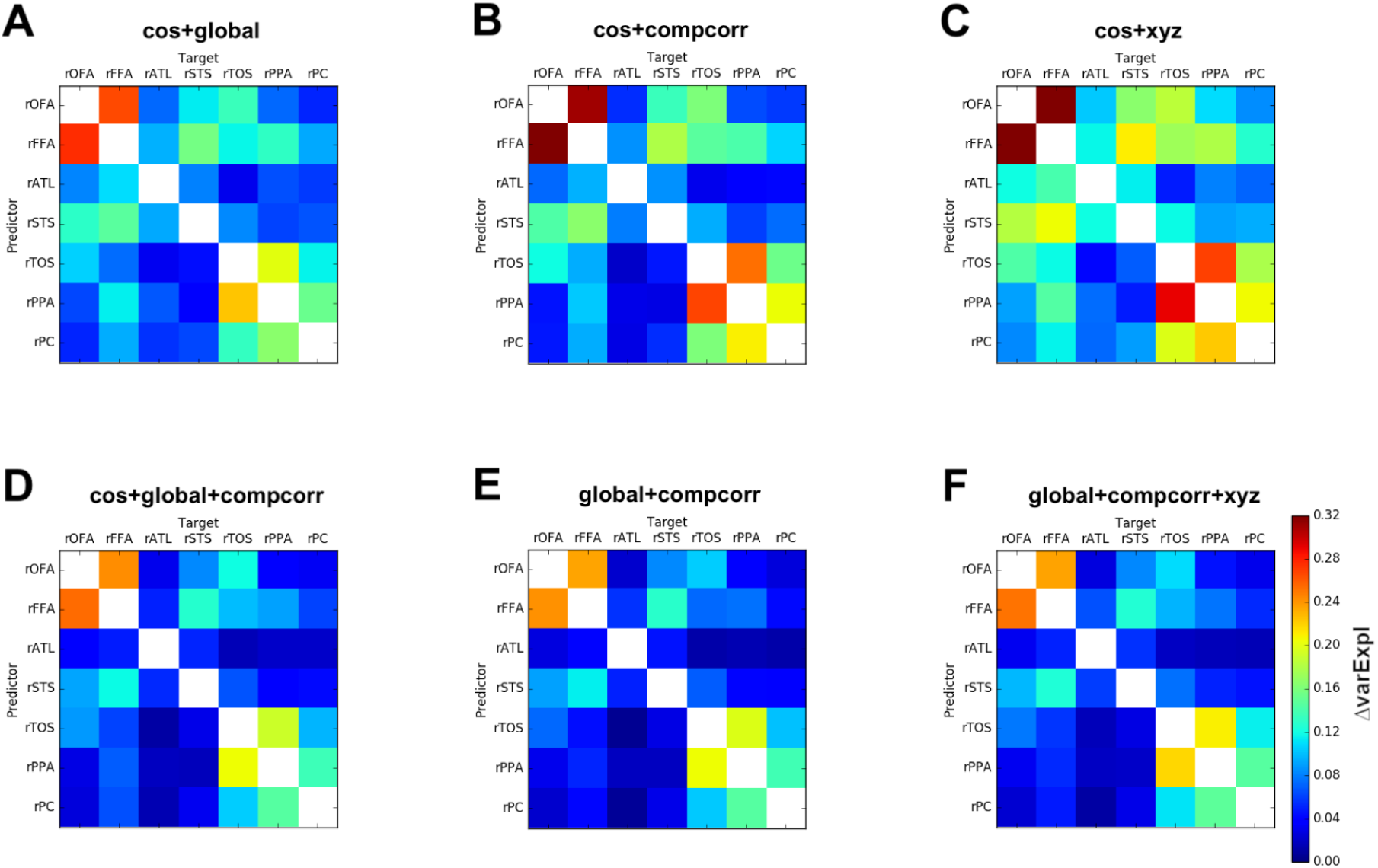
Matrices showing the difference between the variance explained for within-subject MVPD minus the variance explained for iMVPD (Δ varExpl) of A) removal of slow trends followed by removal of global signal, B) removal of slow trends followed by Compcorr, C) removal of slow trends followed by regression of head translation and rotation parameters (motion parameters), D) removal of slow trends followed by removal of global signal followed by Compcorr, E) removal of global signal followed by Compcorr, F) removal of global signal followed by Compcorr followed by regression of head translation and rotation paramaters (motion parmeters).

Adding CompCorr to the removal of slow trends (Figure 5 B) performed better than using the removal of slow trends in isolation (Figure 3 C), however, it did not improve appreciably over using CompCorr alone (Figure 3 E), indicating that the noise eliminated by the removal of slow trends is a subset of the noise captured by CompCorr.

A combination of removal of slow trends and removal of the global signal (Figure 5 A) improved on both methods used in isolation, suggesting that they account for independent sources of noise.

Considering this pattern of results, we expected that using jointly CompCorr and the removal of the global signal would yield optimal results, even as compared to models additionally including removal of slow trends or removal of motion regressors. To test this we computed Δ varExpl matrices for the combination of CompCorr and removal of global signal and found that the combination of these two methods performs better than either CompCorr alone (figure 3 E) or the removal of global signal alone (figure 3 B). Furthermore, by comparing figure 5 (e) with (d) or (f), we found that adding the removal of slow trends or the removal of motion regressors to CompCorr plus the removal of global signal did not further improve Δ varExpl. The combination of CompCorr and removal of global signal achieved parsimonious and effective noise removal.

## 4 Discussion

In this article we have introduced iMVPD, and have shown that it produces similar patterns of statistical dependence between brain regions as standard MVPD (Anzellotti et al., 2017). We have then introduced the difference between the proportion of variance explained within and between participants (hereafter referred to as the ‘discrepancy metric’) and motivated its use as a measure of the effectiveness of denoising methods.

Other methods for studying multivariate interactions exist (i.e. informational connectivity, Coutanche and Thompson-Schill (2013); Crowe et al. (2013), see Anzellotti and Coutanche (2018) for a review), and might have the potential to be extended to perform intersubject analyses. However, the unique combination of 1) attempting to capture most of the variance in each region’s responses, and 2) testing models of connectivity in independent data made MVPD an ideal choice for this study.

Multivariate variants of intersubject connectivity based on canonical correlation analysis have been introduced in previous work (Kriegeskorte, 2015). iMVPD differs substantially from these approaches in that 1) it tests prediction accuracy in independent data, and 2) enables modeling of statistical dependence not only between-subjects within-region, but also between-subjects between-regions.

As predicted, we found that the discrepancy metric was greatest in the absence of denoising (see Figure 3A, F). Different denoising methods varied both in terms of their overall effectiveness (as measured by the average discrepancy metric across all pairs of connections), as well as in terms of the pattern of noise removal across different region pairs. This suggests that different pairs of regions may be differentially affected by distinct sources of noise, and that methods that are very effective at reducing the noise in the interactions between a pair of brain regions may be less effective at reducing noise for other pairs of regions.

Correlation between the discrepancy metric matrices (Figure 4) revealed that removal of the global signal disproportionately affected a different set of connections as compared to removal of slow trends, removal of motion regressor, and CompCorr. This finding suggests that removal of the global signal might target different sources of noise in fMRI data. Specifically, as compared to CompCorr, removal of the global signal disproportionately reduced the discrepancy metric for pairs of regions within a same network (i.e. across pairs of face-selective regions or across pairs of scene-selective regions).

Further studies could investigate the extent to which the different predictors generated by different denoising methods correlate with a variety of physiological measures (i.e. head movement, heart rate, respiration, eye movements) and susceptibility measures to determine whether the noise removed by the global signal has an independent origin.

One possible concern is that removal of the global signal might be subtracting meaningful variation in the fMRI responses. While it is difficult to rule out this possibility entirely, this in itself could not account for the reduction of the discrepancy metric within network: in fact, removal of meaningful signal would reduce both the absolute variance explained within subject as well as the total amount of variance within subject, leaving the proportion of variance explained unchanged (see equation 21).

Testing different combinations of denoising methods revealed that the noise removed by the regression of slow trends and by the regression of motion parameters was mostly a subset of the noise removed by CompCorr. As a consequence, the use of regression of slow trends and regression of motion parameters in addition to CompCorr was redundant. In addition, CompCorr outperformed both of those methods. The effect of motion as measured by rotation and translation parameters on the observed BOLD signal is likely dependent on the anatomical location of voxels, and it is affected by complex phenomena like spin-history effects (in which a voxels move to some extent between slices and spins are no longer excited at regular intervals), and movement across non-uniform regions of the static magnetic field and of the radiofrequency (RF) fields (Liu, 2016). These effects induce nonlinear relationships between motion parameters and the resulting noise, and may be better captured by CompCorr. Across all combinations of denoising methods tested, the discrepancy metric indicated that jointly removing the global signal and applying CompCorr was the optimal denoising pipeline.

Prior studies using simulated data have found that motion artifact causes variable duration of disruptions in signals (Power et al., 2015). Power et al. (2015) also found that proximal correlations are increased by artifact motions more than distal correlations, and that motion regressors have a limited effect in removing motion-related variance, even with voxel-specific regressors or when including a large set of motion regressors. This finding is in line with our result in Figure 3 D that motion parameters have limited efficacy in removing noise. Power et al. (2015) also found that mean white matter or mean ventricle signals are of modest utility as regressors, while fractionation of these signals via ANATICOR (Jo et al., 2010) or aCompCorr (Muschelli et al., 2014) or other methods may provide additional benefit. With iMVPD and the discrepancy metric we find that aCompCorr using multiple dimensions extracted from white matter and the cerebrospinal fluid (Figure 3 E) provides considerable effectiveness in removing noise as compared with motion regressors only (Figure 3 D). Power et al. (2015) pointed out that regressing the global signal is an effective processing step. In our findings (Figure 3 B), removal of global signal is disproportionately effective in removing noise within-network (among face-selective region pairs or among scene-selective region pairs), and less effective in removing noise from between-network region pairs.

While these findings advance our understanding of denoising methods for fMRI, they are nonetheless affected by some limitations. First, optimal denoising methods are expected to minimize the discrepancy metric, however, it is worth noting that even perfect denoising would not reduce the discrepancy metric to zero. This is because individual differences between participants are expected to reduce iMVPD as compared to standard within-subject MVPD: the discrepancy metric is a result of both noise and individual differences. The discrepancy metric is a useful tool to compare the performance of different denoising methods in real fMRI data. However, it is important to note that while it provides a relative measure that can be used to compare different denoising methods, it is not an absolute measure of denoising. Therefore, the discrepancy metric cannot be used to determine exactly how much noise is left in the data. Second, our study was necessarily restricted to a subset of the vast set of denoising approaches that have been developed for fMRI. For example, we did not consider interpolation (Power et al., 2014), ANATICOR (Jo et al., 2010), group-level covariates and partial correlations. Future studies could use the discrepancy metric to test additional denoising approaches that have not been included in the present investigation. Another effective strategy we did not test in this study consists in censoring motion-contaminated data. In the context of intersubject analysis, censoring would require removing all timepoints that show excessive motion in either of two subjects, further reducing the available data. Despite we did not test this approach in the present study, it can be applied in conjunction with the removal of global signal plus CompCorr approach we recommend. Finally, it will be important to extend the present results to other datasets employing different tasks and studying different brain regions to test the generality of the present findings.

In this study, we have used iMVPD as an instrument to define a discrepancy metric for denoising methods in order to assess their performance on ‘real’ (non-simulated) fMRI data. Among the methods tested, a combination of CompCorr and removal of the global signal was the most effective. Beyond the scope of this study, iMVPD could be used for a variety of applications. In ongoing work, we are using iMVPD to study individual differences across participants. Another potential area of application for iMVPD is the analysis of imaging modalities in which it is challenging to acquire data simultaneously from multiple regions, such as electrocorticography (ECoG) or primate electrophysiology. More generally, iMVPD can be applied as a conservative approach to study multivariate interactions between brain regions.

## Acknowledgments

We would like to thank the researchers who contributed to the studyforrest project (Hanke et al., 2016; Sengupta et al., 2016) for sharing their data, and the developers of fmriprep (Esteban et al., 2018) for their assistance with the fmriprep preprocessing pipeline.

